# Expanding gene regulatory networks from transcriptome data through graphical modeling with heterogeneous priors

**DOI:** 10.64898/2026.06.12.731835

**Authors:** Toshiya Kokaji, Kenta T. Suzuki, Katsuyuki Kunida, Yuichi Sakumura

## Abstract

Gene regulatory network inference is widely used to reconstruct large-scale networks and identify functional genes from transcriptome data. Meanwhile, in many biological fields, core regulatory genes have been extensively studied, leading to the establishment of small-scale gene regulatory networks, and novel genes connected to these networks remain to be identified. However, methods for expanding existing gene networks by identifying novel regulatory interactions, rather than reconstructing the entire network, are not well established. Here, we propose a method for gene network expansion that incorporates known regulatory relationships and evaluates each candidate gene individually to infer its regulatory connections to the existing network. Using simulated datasets from the DREAM4 benchmark and the PRECISE-1K experimental dataset, our method outperformed conventional methods by incorporating prior knowledge. In particular, it improved the ability to distinguish true regulatory interactions from indirect associations arising from strong correlations among genes in the existing network. The method also showed strong performance for interactions involving genes with high outdegree or centrality. Furthermore, it maintained stable performance as the size of the existing network increased and was robust to noise in prior information. These results demonstrate that our method provides an effective framework for expanding existing gene regulatory networks by leveraging prior knowledge.

## Introduction

Gene expression is orchestrated through complex regulatory interactions forming networks. Statistical inference of gene regulatory networks from large-scale gene expression data is a useful approach for identifying novel functional genes^1–3^. Since the advent of microarray and RNA-seq technologies, which enable large-scale expression profiling, numerous network inference methods have been developed. Their performance has been extensively evaluated using both simulated and experimental datasets for which ground-truth networks are available^4–6^. In addition, methods have been proposed that integrate additional regulatory evidence, such as ChIP-seq profiles and transcription factor binding motifs, to infer transcription factor (TF)–target relationships^7–9^.

Most existing network inference methods aim to reconstruct networks over all measured genes. Meanwhile, numerous experimental studies in molecular biology have individually characterized regulatory relationships among genes, and continue to identify novel genes to be incorporated into these existing networks. When identifying novel regulatory genes for an existing network, genome-wide inference introduces an unnecessarily broad scope. Specifically, it redundantly infers interactions entirely outside the network and established interactions within the network. A more tailored framework would instead infer regulatory relationships between genes inside and outside the network while keeping established regulatory relationships fixed; however, such an approach remains poorly established.

Known regulatory relationships can be incorporated as prior information in network inference. Many approaches adopt weighting schemes for interactions supported by prior information. For example, weighted graphical LASSO (wgLASSO) facilitates the inference of regulatory relationships by reducing the L1 penalty for interactions with prior support^8,10^. Other approaches exclude interactions without prior information from the estimation process^9^. While these strategies prioritize known relationships, they do not enforce them as fixed constraints. Consequently, such approaches are well suited for large-scale, noisy information such as ChIP-seq profiles^7^; however, they are less effective for high-confidence, experimentally validated regulatory relationships. Furthermore, they cannot incorporate information about the sign of regulatory effects—activation or repression.

Gene regulatory networks are generally sparse and exhibit heterogeneous connectivity, with a small number of hub regulators controlling many target genes^11,12^, and variable selection has therefore been widely applied in network inference. The graphical LASSO (gLASSO) enables simultaneous variable selection and parameter estimation within a single optimization framework. However, because all coefficients are estimated simultaneously, highly correlated explanatory variables often compete to explain shared variation, leading to mutual interference that can obscure their individual effects. In high-dimensional gene expression data with complex correlation structures, L1 regularization may further attenuate true regulatory relationships. This motivates the need for a framework that allows for the independent evaluation of candidate interactions. In forward wrapper approaches, explanatory variables are tested sequentially, enabling the independent assessment of each candidate variable relative to the current model, thereby preventing the mutual interference caused by simultaneous estimation. Although this approach is computationally intensive due to the large number of models evaluated, it effectively decouples the evaluation of unselected candidates.

To address these issues, we developed a novel gene regulatory network inference method: Graphical Network Assembly and Inference using Stepwise Testing with Optimization (gNAISTO). This method was developed by extending our previously developed algorithm for application to precision matrix estimation^13^. Given an expression dataset and a gene network, gNAISTO identifies novel regulatory interactions involving genes in the existing network by incorporating a per-gene inference framework and customized prior distributions. We evaluated the performance of gNAISTO using both simulated and experimental datasets. The results demonstrate that gNAISTO achieves superior inference performance in both settings, highlighting the efficacy of leveraging prior information to identify novel regulatory relationships. Furthermore, gNAISTO outperforms existing methods as the extent of prior information expands and remains robust to noise in the prior information. Finally, analysis of the inferred edges reveals that gNAISTO achieves particularly high performance for interactions involving highly connected nodes.

## Results

### Conceptual overview of network inference

To identify novel genes that connect to an existing gene network using an expression dataset, we developed a framework: Graphical Network Assembly and Inference using Stepwise Testing with Optimization (gNAISTO; Fig. 1A). In this framework, we define the provided network as “known network”, genes within it as “known genes”, and all remaining genes as “unknown genes”. Unlike conventional network inference approaches, gNAISTO does not aim to reconstruct the entire network; instead, it evaluates the connectivity of unknown genes to the known network (Fig. 1B). Accordingly, regulatory edges among unknown genes are not estimated, and inference is focused on edges involving known genes (Fig. 1B, gray region). We assess performance by comparing the inferred edges between known and unknown genes with ground-truth data.

**Fig. 1.**
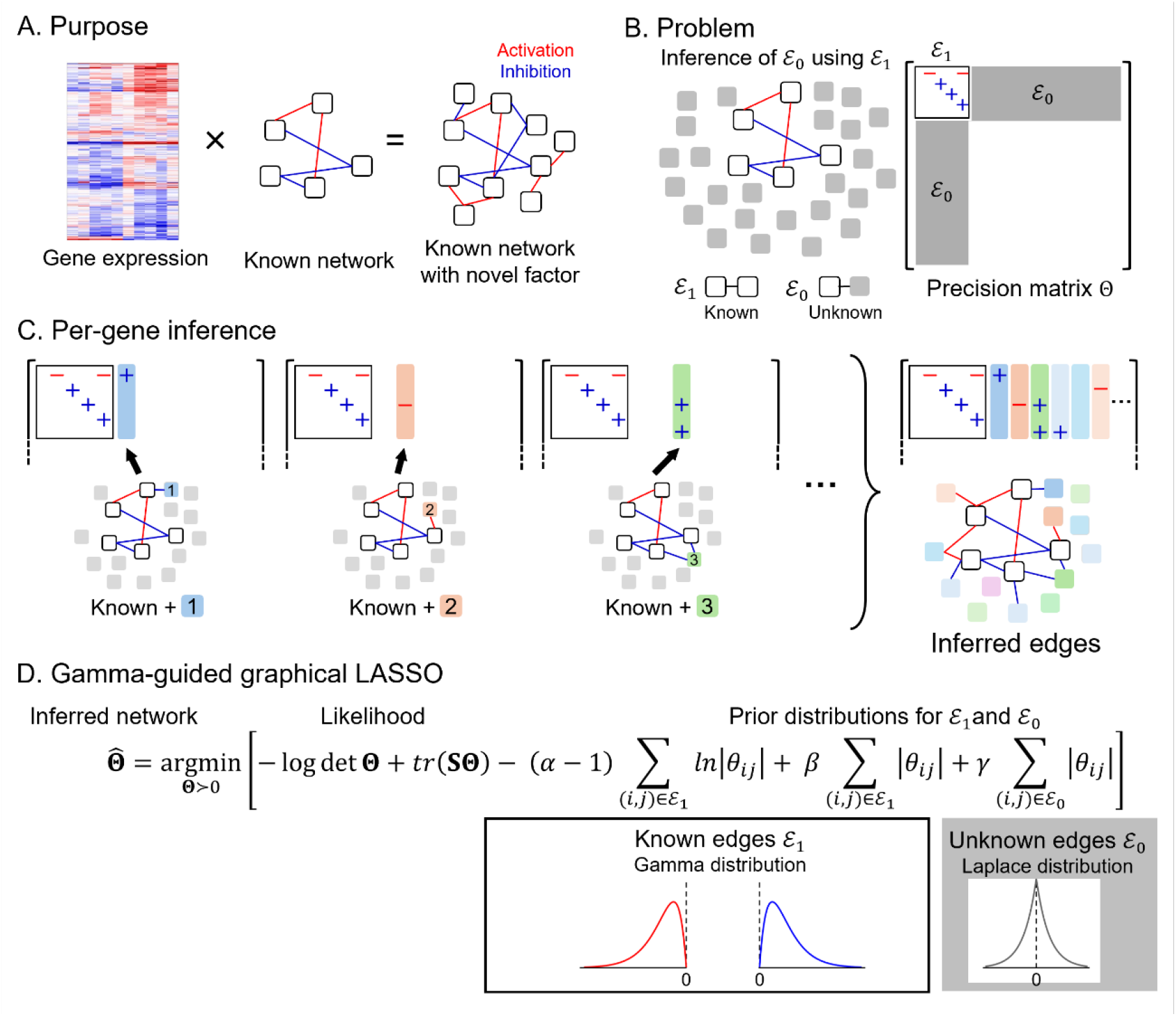
Overview of the proposed network inference framework (gNAISTO). (A) Our method identifies novel genes connected to a known gene network by integrating gene expression data with prior network information. (B) The method estimates regulatory edges by optimizing the precision matrix **Θ**, which represents the gene network structure. Using prior information on known edges (ℰ_1_), edges between genes in the known network and unknown genes (ℰ_0_) are inferred. Note that edges are defined by the partial correlation coefficient 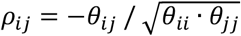, meaning that a negative *θ*_*ii*_ corresponds to an activating edge, while a positive *θ*_*ii*_ represents an inhibitory edge. (C) Edges between the known network and unknown genes are obtained from networks inferred by adding each unknown gene individually to the known network. (D) In the optimization, different regularization terms are applied to known edges (ℰ_1_) and unknown edges (ℰ_0_). Unknown edges are penalized using an L1 term corresponding to a Laplace prior. For known edges, a gamma prior is imposed via a combination of an L1 penalty and a logarithmic term. Note that edges are defined by the partial correlation coefficient 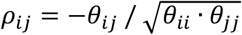, meaning that a negative *θ*_*ii*_ corresponds to an activating edge, while a positive *θ*_*ii*_ represents an inhibitory edge.

To enhance the identification of novel genes, we introduce a per-gene inference framework for unknown genes and a customized representation of known regulatory relationships (Fig. 1, C and D). Rather than incorporating all unknown genes simultaneously, our strategy evaluates the regulatory connections of each unknown gene individually (Fig. 1C). In gene regulatory networks, highly connected hub genes are often associated with clusters of correlated genes. In simultaneous optimization frameworks, correlated candidate genes often interfere with one another, making their individual regulatory contributions difficult to disentangle and thereby obscuring true regulatory edges. In contrast, our per-gene inference framework allows the contribution of each unknown gene to be assessed independently, reducing interference from other unknown genes. While this procedure increases computational cost for separate optimization, the independent nature of these tasks allows for efficient parallelization, ensuring the overall cost remains tractable. In addition, the solutions obtained for each gene are reused as initial values for subsequent optimizations to accelerate convergence (see Methods).

To explicitly incorporate edges from the known network, we extend gLASSO by introducing an additional term corresponding to a gamma distribution (Fig. 1D). The gLASSO formulates network inference as an optimization problem over the precision matrix **Θ**, where each off-diagonal element *θ*_*ii*_ represents the regulatory interaction between genes *i* and *j*^14^. The objective function can be interpreted as the log-posterior in a Bayesian context. The data-fitting term, reflecting the discrepancy between the sample covariance matrix **S** and **Θ**, corresponds to the log-likelihood, while the L1 penalty represents the log-prior under a Laplace distribution over regulatory edges (Fig. 1D, gray). Because the Laplace distribution is sharply peaked at zero, it promotes sparsity by shrinking coefficients toward zero. To account for known regulatory interactions, we employ a gamma distribution as a prior, distinct from the Laplace prior used for unknown ones. This formulation promotes nonzero values for known edges, preventing them from shrinkage toward zero (Fig. 1D, red and blue). While wgLASSO also incorporates prior information, it employs varying Laplace priors and is therefore better suited for large-scale, potentially noisy screening data. By adjusting the hyperparameters of the gamma distribution, the strength of the constraints on known edges can be finely tuned, ranging from fixing coefficients near the prior mode to recovering standard gLASSO behavior (Fig. S1).

The resulting optimization problem can be decomposed into column-wise subproblems, analogous to the standard gLASSO. Each subproblem is formulated as a LASSO regression integrated with a gamma-based prior; importantly, the problem remains convex, enabling efficient optimization via proximal gradient methods. We also developed a tuning strategy that automates the selection of the hyperparameters (*α, β, γ*) by leveraging the known network. Specifically, a subset of the known regulatory edges is masked during tuning, and the optimal hyperparameter configuration is selected based on its performance in recovering these masked edges. This strategy ensures the automated selection of appropriate regularization strengths, adapted to the specific reliability of prior information (see Methods for details).

### Performance of gNAISTO across simulated and experimental datasets

To evaluate the performance of the proposed method for gene network inference, we applied gNAISTO to both simulated and experimental datasets (Fig. 2). For each dataset, 20 genes were randomly selected from the ground-truth network as the known network in 50 trials, and regulatory edges between these known genes and the remaining unknown genes were inferred. We employed the area under the precision–recall curve (AUPRC) as the performance metric, as it emphasizes the precision of top-ranked predictions for identifying candidate genes. During AUPRC calculation, we accounted for the regulatory sign (activation or repression) but not the edge directionality (see Methods). The performance of gNAISTO was benchmarked against two methods that incorporate prior information, wgLASSO and semi-supervised SVM (S3VM), as well as GENIE3, a widely used method that does not use prior information^10,15,16^.

**Fig. 2.**
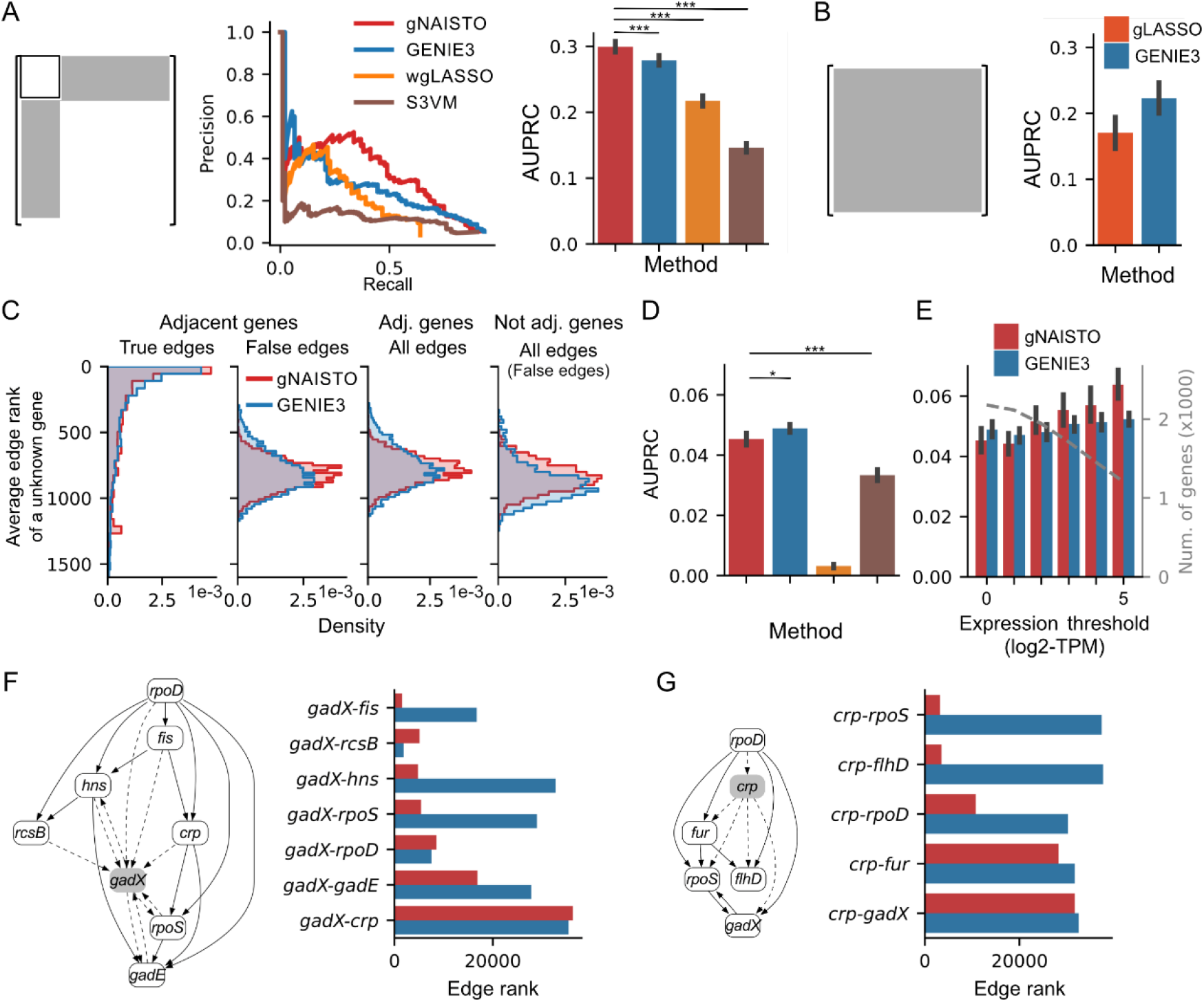
Performance across methods in simulated and experimental datasets. (A) Precision–recall curves and corresponding AUPRC values for the inference of regulatory edges between known and unknown genes using simulated datasets from the DREAM4 benchmark. Known networks were defined by randomly selecting 20 genes from the ground-truth networks; this subnetwork size was used for all subsequent panels unless otherwise specified. Bar plots represent the mean AUPRC, with error bars indicating the standard error (*SE*) across 50 trials (5 datasets × 10 random subnetworks). Statistical significance was assessed using the Wilcoxon signed-rank test (\**P* < 0.05, \*\**P* < 0.01, \*\*\**P* < 0.001). (B) Mean AUPRC for whole-network inference on the DREAM4 datasets, provided as a baseline comparison. For gLASSO, the regularization parameter was selected to maximize the AUPRC. (C) Distributions of the average rank of inferred edges from a single unknown gene to the known genes using gNAISTO (red) and GENIE3 (blue). The left panels show the average ranks of true and false regulatory edges between known genes and adjacent unknown genes. The right panels show the average ranks of all inferred edges between known genes and either adjacent or non-adjacent unknown genes. (D) Evaluation on the PRECISE-1K experimental dataset. Error bars indicate the *SE* across 50 random subnetworks. Results for wgLASSO are excluded for cases where optimization failed to converge or hyperparameter tuning was unsuccessful (see Methods). (E) Mean AUPRC on the PRECISE-1K dataset under various expression thresholds. The number of genes is also shown (gray line). Genes with no connections in the network were excluded. (F, G) Representative trials showing known networks associated with *gadX* (F) or *crp* (G) and the edge ranks of associated regulatory interactions.

We first compared inference performance using simulated datasets from the DREAM4 benchmark, which comprises five gene regulatory networks derived from *E. coli* and *S. cerevisiae*^4,17,18^. In this benchmark, gNAISTO achieved the highest average AUPRC (0.30; Fig. 2A). Notably, the standard graphical LASSO without explicit prior information exhibited substantially lower performance than GENIE3 (Fig. 2B), indicating that the incorporation of known network information markedly improves the inference of regulatory relationships. Furthermore, gNAISTO showed a significant improvement in AUPRC compared with wgLASSO, largely attributable to the per-gene inference framework (Fig. S2). The relatively low performance of S3VM may be limited by the small size of the known network available for training, as this approach typically requires larger training sets to optimize its decision boundaries.

Genes within the known network are often correlated, and such correlations can propagate to genes adjacent to the network, leading to spurious associations. Because gNAISTO estimates partial correlations while incorporating prior information from the known network, it can effectively mitigate these indirect correlations. To examine this effect, we evaluated the rank distributions of false edges connecting the known network to adjacent genes (Fig. 2C). In gNAISTO, the rank distributions of false edges were similar regardless of whether they involved adjacent or non-adjacent genes. In contrast, in GENIE3, false edges involving adjacent genes tended to receive higher ranks, indicating a higher rate of false positives arising from indirect paths. Since the rank distributions of true edges were comparable between the two methods, the superior AUPRC of gNAISTO is primarily attributable to its improved ability to distinguish true interactions from false edges involving adjacent genes.

We next evaluated inference performance using large-scale experimental data (Fig. 2D). We utilized the PRECISE-1K dataset^19^, which comprises over 1,000 *E. coli* RNA-seq samples covering 4,257 genes, with partially annotated regulatory relationships from RegulonDB^20^. Consistent with previous reports^5,6^, overall AUPRC values were lower for experimental data than for simulated datasets. Although gNAISTO outperformed wgLASSO and S3VM, its performance was comparable to that of GENIE3. Because lowly expressed genes are associated with high noise levels, we performed our analysis under various expression thresholds (Fig. 2E). When restricting the analysis to genes with average TPM above 2^2^, gNAISTO consistently outperformed GENIE3 across all tested thresholds, achieving its highest average AUPRC of 0.065 at a TPM thresholds of 2^5^. In some trials, we also observed substantially improved edge rankings compared with GENIE3 for regulatory edges involving *gadX* (Fig. 2F), a regulator of acid resistance genes^21,22^, and *crp* (Fig. 2G), a global regulator of carbon metabolism^23,24^. These results demonstrate that gNAISTO consistently enhances predictive performance across both simulated and experimental datasets by leveraging prior network information.

### Factors influencing gNAISTO performance

To characterize the input conditions under which gNAISTO performs optimally, we systematically evaluated its performance across various conditions using the DREAM4 simulated dataset. First, we assessed robustness to measurement noise by introducing varying noise levels into the expression data (Fig. 3A). While all methods showed lower AUPRC as noise increased, this effect was most pronounced for gNAISTO. This behavior can be attributed to its reliance on linear regression, which is generally more sensitive to noise and outliers than tree-based methods^25,26^. These results confirm that high-quality expression data enhances the performance of gNAISTO, consistent with our earlier findings from the *E. coli* dataset analysis (Fig. 2E). Next, we evaluated robustness to prior network noise by introducing spurious edges into the known networks (Fig. 3B). Notably, gNAISTO maintained its performance even as the number of false edges increased, achieving an AUPRC comparable to GENIE3 even when the number of false edges was set to twice the number of true edges. This result suggests that gNAISTO flexibly adjusts the gamma prior to suppress coefficients of spurious edges while preserving those of true interactions. We further examined the impact of the known network size by varying the number of known genes from 10 to 25 (Fig. 3C). Interestingly, the AUPRC of gNAISTO remained nearly constant despite the increasing network size. In contrast, methods such as GENIE3 exhibited a declining trend in AUPRC. This decline is likely attributable to the decreasing baseline AUPRC, as the positive rate among candidate edges decreases with increasing network size (gray line). Consequently, gNAISTO appears to effectively leverage the additional structural information provided by the larger known network.

**Fig. 3.**
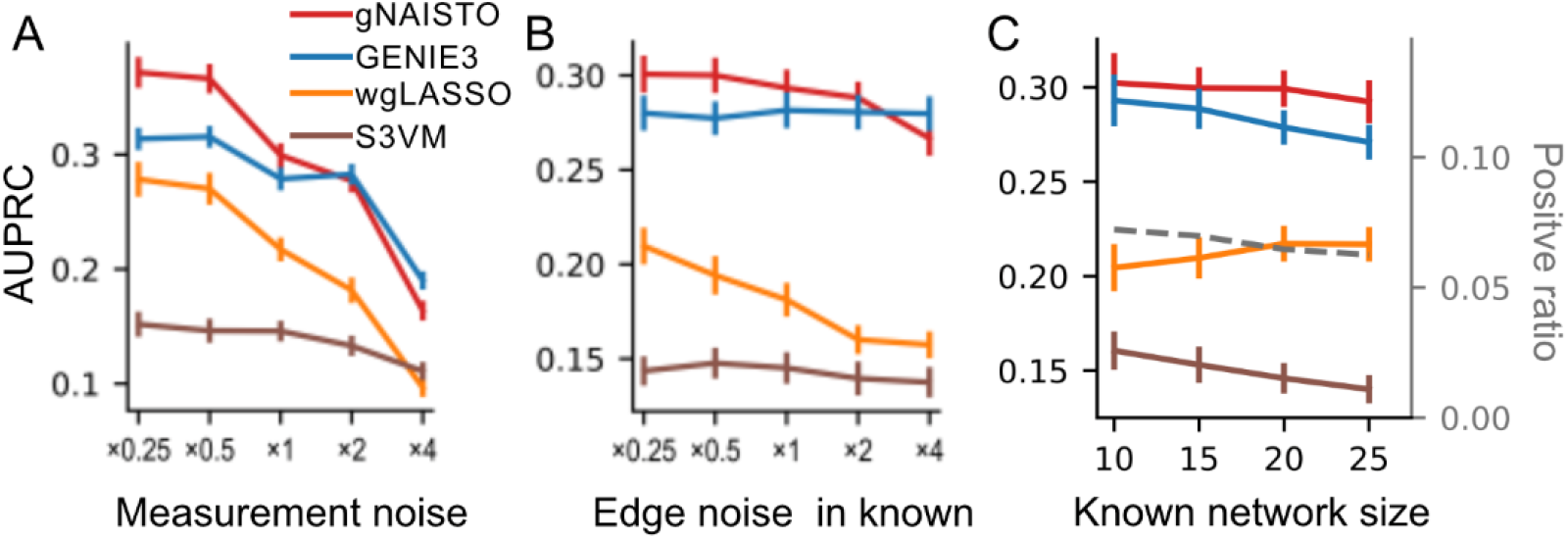
Effect of input data characteristics on gNAISTO performance. (A–C) AUPRC under varying input conditions using DREAM4 dataset. (A) Effect of measurement noise in expression data. (B) Effect of noise in prior network, represented by the number of spurious edges added to the known network. (C) Effect of the size of the known network (number of nodes). Measurement noise is scaled relative to the original DREAM4 data (×1). Edge noise is defined relative to the number of true edges in the known network (×1). Error bars indicate the *SE* across 5 datasets × 10 random subnetworks. In (C), the positive rate of candidate edges (corresponding to the baseline AUPRC under random prediction) is also shown (gray line).

To characterize the types of regulatory edges preferentially inferred by gNAISTO, we analyzed features of true regulatory edges that showed rank improvements relative to GENIE3 in the DREAM4 dataset (Fig. 4A). We focused on edges from unknown genes to known genes in the ground-truth network, representing the identification of true regulatory factors. Edge features were derived from the features of both endpoint genes (unknown and known genes) as well as their combined metrics. We then computed Spearman’s rank correlations between these features and the changes in edge ranking. The results revealed that features of known genes (white) were more strongly associated with ranking improvements than those of unknown genes (gray). In particular, edges involving known genes with high outdegree exhibited higher ranks in gNAISTO (Fig. 4B). A high outdegree implies that many unknown genes are correlated with the known gene, where the per-gene inference framework may mitigate mutual interference among correlated unknown genes. In contrast, edges targeting known genes with high indegree were more difficult to infer (Fig. 4C), especially when the known gene receives inputs from multiple unknown genes (Fig. 4D). In such cases, the per-gene inference framework excluded other relevant predictors that should ideally be considered jointly, leading to a lack of necessary covariates to explain the variance of the target gene. This limitation may be alleviated by extending the framework to small groups of unknown genes (e.g., pairs or triplets). Finally, the positive correlation with eigenvector centrality (Fig. 4E) indicates that gNAISTO performs particularly well in inferring regulatory edges involving hub genes.

**Fig. 4.**
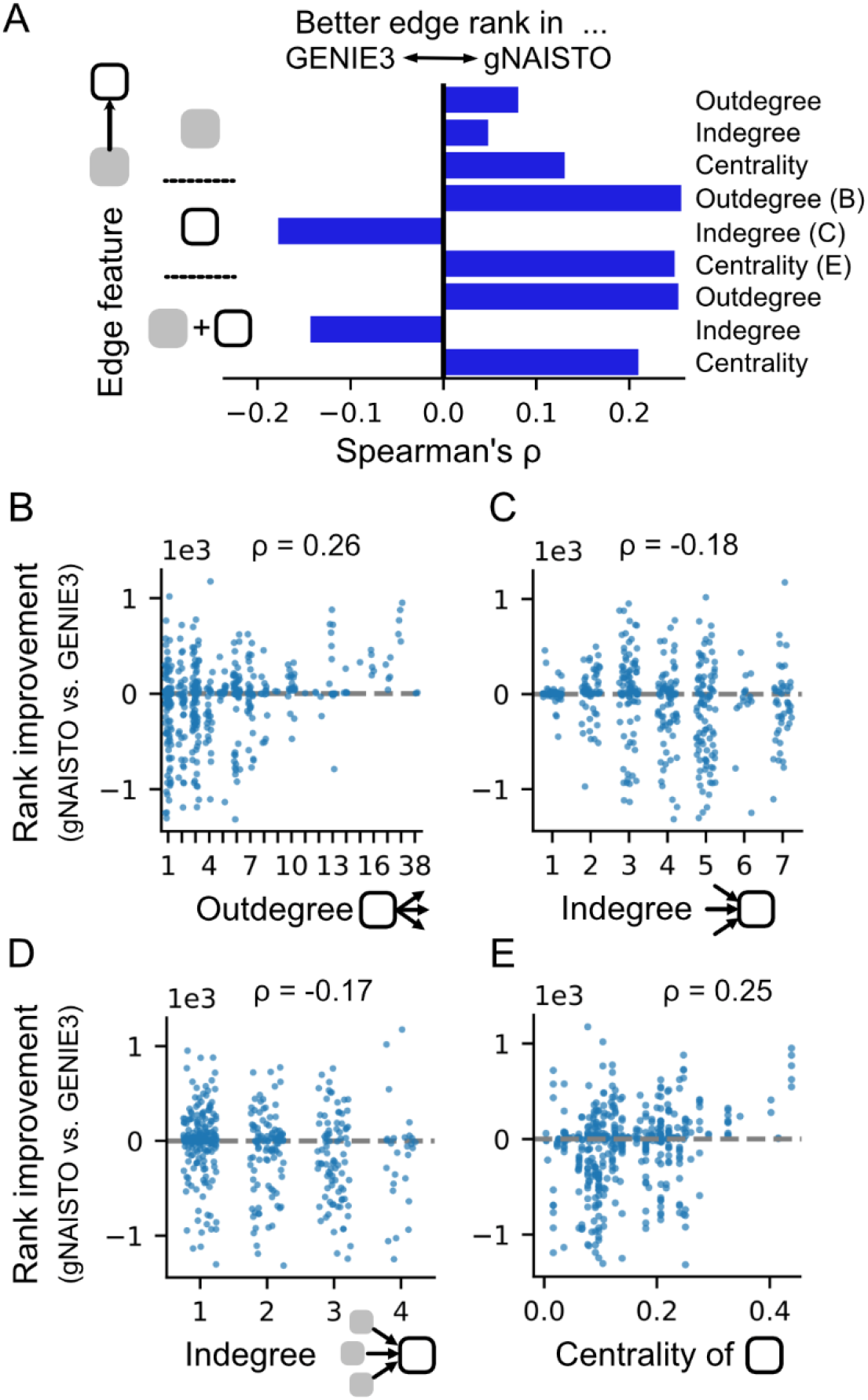
Effect of edge characteristics on gNAISTO performance. (A) Spearman’s correlation coefficients (ρ) between edge features and rank improvement in gNAISTO relative to GENIE3. True edges from 50 trials on the DREAM4 dataset (as in Fig. 2A) were used for the analysis. The analysis focuses on true edges from unknown genes to known genes. Edge features are defined using features of the endpoint nodes—unknown (gray) and known (white)—including outdegree, indegree, eigenvector centrality, and their sums. (B–E) Rank improvement in gNAISTO relative to GENIE3 versus edge features. Features include outdegree of known nodes (B), indegree of known nodes (C), indegree from unknown to known nodes (D), and eigenvector centrality of known nodes (E).

## Discussion

In this study, we proposed gNAISTO, a method for identifying novel regulatory relationships linked to a known gene network, and demonstrated its high inference performance in both simulated and experimental datasets. Our method extends the conventional wgLASSO framework by introducing a gamma-distribution prior for known regulatory interactions and regulatory edge inference for each candidate gene. Bayesian frameworks have been widely applied to gene regulatory network inference^27–29^. In this study, we formulate gene network inference incorporating known regulatory information as a maximum a posteriori estimation problem for the precision matrix, introducing gamma distributions as priors. To solve this problem, we developed an optimization algorithm that decomposes the objective into subproblems using block coordinate descent and solves each subproblem using a proximal gradient method. We found that gNAISTO effectively distinguishes true regulatory interactions from false positives among edges involving genes adjacent to the known network, suggesting that partial correlation estimation under the constraint of the known network structure helps filter out conditionally independent relationships. In addition, gNAISTO showed improved performance in identifying novel regulatory relationships for hub-like nodes with high outdegree and centrality, indicating that the per-gene inference framework mitigates mutual interference among highly correlated genes during coefficient estimation. We also found that gNAISTO was robust to noise in prior network information but sensitive to measurement noise in expression data, highlighting the importance of expression data quality, as observed in the PRECISE-1K analysis. As expected, gNAISTO showed better performance over the other methods when larger known networks were provided. Therefore, we recommend expanding the known network using regulatory interactions from curated databases when available^20,30–32^.

The central concept underlying our framework is the existence of a known gene network. In transcriptomic studies, it is common that a subset of genes of interest and their regulatory relationships have already been partially characterized through prior experimental efforts. First, by focusing on network expansion, the inference can be restricted to regulatory relationships involving genes within the known network. The per-gene inference framework is specifically designed for this context, as it does not estimate interactions among unknown genes. This design contributed to improved performance compared with methods such as wgLASSO, which require full-network estimation. Notably, such restriction of the inference space can be applied to other methods; for instance, limiting the target variables in GENIE3 to the known genes substantially reduces computational cost. Second, the known gene network serves as a set of partially labeled edges. In gene regulatory network inference, hyperparameters are typically tuned using indirect criteria, such as model fit to expression data or the stability of inferred edges under subsampling^8,33^. In contrast, our framework enables direct evaluation of edge inference by leveraging the known gene network as labeled regulatory edges. Specifically, by masking known edges and assessing their recovery, hyperparameters can be optimized based on the accuracy of edge inference itself.

The incorporation of transcription factor (TF) information, including ChIP-seq profiles and binding motifs, has been widely reported to improve the accuracy of gene regulatory network inference^7–9^, and our framework can readily leverage such information. For example, weighting schemes using TF priors can be incorporated into the gNAISTO objective function. In this work, we focused on gene expression data; however, gNAISTO can also be applied to inferred TF activities derived from TF information^7,34,35^. In TF-based regulatory inference, the partial correlation estimation in gNAISTO may help improve robustness when inferred TF activities are correlated due to overlapping binding profiles or similar motifs. A common limitation of expression-based network inference methods, including ours, is the difficulty of determining the regulatory directionality^16,36^. Restricting the known network to transcription factors may help constrain the directionality of inferred regulatory interactions.

In this study, we systematically characterized the performance of gNAISTO and identified several practical considerations. As with many gene regulatory network inference methods, gNAISTO is most effective when applied to expression datasets with a sufficient sample size to enable reliable estimation of regulatory relationships^1,4,37^. From this perspective, single-cell RNA-seq (scRNA-seq) data represent a promising resource^3,38,39^. However, because gNAISTO is sensitive to measurement noise (Fig. 3A), careful preprocessing—such as gene selection and cell clustering—is likely to be critical for reducing noise and improving inference accuracy in scRNA-seq analyses. We also found that the inference of regulatory edges targeting genes with multiple uncharacterized regulators is challenging (Fig. 4D). This limitation likely arises from the per-gene inference framework, which simplifies the model by excluding potential predictors that might ideally be considered jointly. While simultaneous evaluation of multiple genes may alleviate this issue by allowing for more appropriate model complexity, this approach also leads to a combinatorial growth in computational cost (e.g., 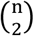 gene pairs), requiring additional strategies such as gene filtering. Overall, gNAISTO provides a practical framework for screening novel regulatory interactions that aligns with standard molecular biology workflows, thereby facilitating hypothesis generation for experimental validation. Future efforts will focus on developing user-friendly implementations tailored for experimental biologists to further broaden the applicability of the method.

## Methods

### gNAISTO

gNAISTO is a framework designed to identify novel genes connected to a known gene network by leveraging gene expression data together with prior regulatory information. The framework consists of two main components: a per-gene inference framework and a gamma-guided graphical LASSO.

### Per-gene inference framework

To infer regulatory edges between unknown genes and a known gene network, we estimate regulatory interactions by adding one unknown gene at a time to the known network. Formally, given a known network of *N* genes and *M* unknown genes, gamma-guided graphical LASSO (see below) is applied *M* times, each time using a set of *N* + 1 genes consisting of the known network and a single unknown gene. For computational efficiency, inference for each unknown gene is parallelized (number of cores = 8). The batch size for parallel processing is gradually increased (8, 16, 32, 64, …). The optimal solutions obtained in each batch are reused as candidate initial values for subsequent batches (see below).

### Gamma-guided graphical LASSO

To estimate a sparse precision matrix representing conditional dependencies among genes while incorporating known activation and repression relationships, we developed gamma-guided graphical LASSO. This method extends the conventional graphical LASSO (gLASSO), which estimates conditional dependencies based on covariance structure, by incorporating known network information as statistical priors.

Given a sample covariance matrix **S** ∈ ℝ^*p*×*p*^, gNAISTO estimates the precision matrix **Θ** by solving the following optimization problem:

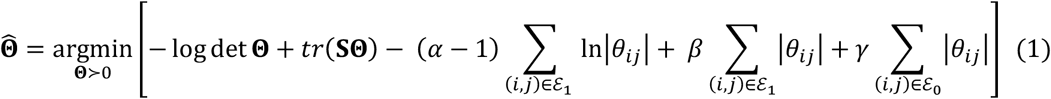

Alternatively,

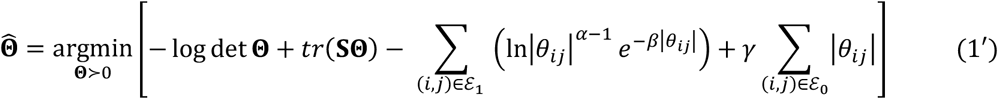

Here, **Θ** denotes the precision matrix, and *θ*_*ii*_ represents its (*i, j*)-th element. ℰ_1_ denotes the set of edges with prior information, where activation or repression relationships are known, while ℰ_0_ denotes the set of edges without prior information. Note that edges within the known network that lack prior information are assigned to ℰ_0_ and are not fixed to zero. The parameters *α* and *β* correspond to the shape and rate parameter of the gamma distribution, respectively. The parameter *γ* corresponds to the regularization parameter in conventional gLASSO.

From the estimated precision matrix **Θ**, the partial correlation matrix **R** = [*ρ*_*ii*_] is computed as:

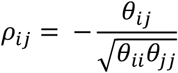

The partial correlation matrix *R* represents conditional dependencies between variables, and *ρ*_*ii*_ is used as the estimated score for regulatory interactions. Due to the sign inversion between the precision matrix and the partial correlation matrix, activation corresponds to a negative gamma prior, whereas repression corresponds to a positive gamma prior. In the objective function in (Eq. 1), the absolute value *θ*_*ii*_ is used, allowing the log-likelihood contributions of these two gamma priors to be combined into a single term.

The optimization of Eq. (1) is performed using block coordinate descent (BCD), as in gLASSO^14,40^. The precision matrix **Θ** and covariance matrix **S** are partitioned with respect to column *j* as follows:

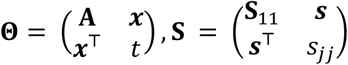

Here **A** ∈ ℝ^*p*−1×*p*−1^, ***x*** ∈ ℝ^*p*−1^, and *t* ∈ ℝ denote the corresponding block components. Using standard block matrix identities, we obtain:

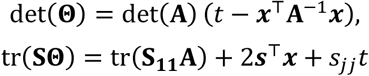

Substituting these expressions into the objective function in Eq. (1) and ignoring constant terms with respect to *x* and *t* the subproblem for column *j* can be written as:

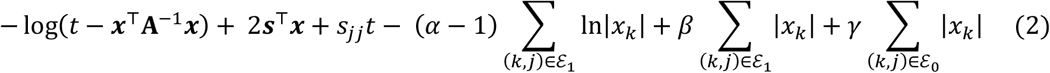

Differentiating with respect to *t*, we obtain:

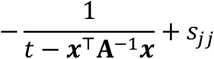

Setting this derivative to zero yields the optimal value:

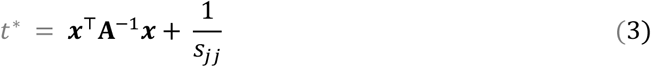

Substituting Eq. (3) into Eq. (2), the optimization problem with respect to *x* is:

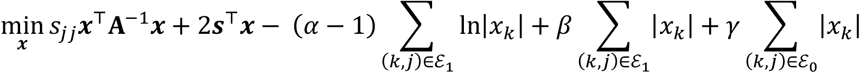

Rescaling variables as *x*^′^ = *s*_*ii*_*x*, the problem becomes:

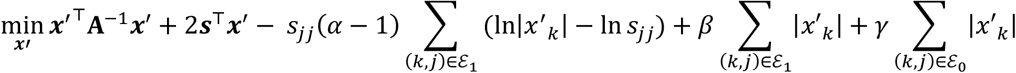

Dropping constant terms involving ln *s*_*ii*_, we obtain the following extended LASSO subproblem:

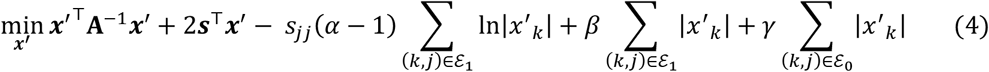

The regression coefficients obtained by solving this subproblem are used to update the corresponding column (and row) of the precision matrix. Note that, compared with Eq. (1), the coefficient of the log-term becomes *s*_*jj*_(*α* − 1), rather than (*α* − 1). All columns are updated iteratively until the relative change in the objective function falls below 10^−4^.

To accelerate optimization, the initial value of the precision matrix **Θ** was selected from multiple candidates as the one yielding the lowest value of the objective function in (Eq. 1). The candidate initial values included (i) the raw precision matrix obtained directly from the sample covariance matrix, (ii) the precision matrix estimated by gLASSO with the same regularization parameter *γ*, (iii) matrices derived from (i) and (ii) by replacing the entries corresponding to known regulatory edges ℰ_1_ with the mode of the gamma distribution, and (iv) previously obtained optimal solutions (see above). In addition, when the sign of an entry corresponding to ℰ_1_ was inconsistent with the known regulatory sign (activation or repression), modified matrices were generated as additional candidates by replacing the corresponding entries with either the mode of the gamma distribution or a small value (10^-10^) with the appropriate sign. Due to the sign inversion between the precision matrix and the partial correlation matrix, entries corresponding to activation are assigned negative values, whereas those corresponding to repression are assigned positive values. Non–positive definite matrices were excluded from the candidate set.

The extended LASSO subproblem (Eq. 4) is convex. Because the objective function consists of a smooth differentiable term, including quadratic and logarithmic components, and non-smooth L1 penalty terms, we employed an iterative optimization scheme using proximal gradient methods. Specifically, the objective function was decomposed into a smooth term

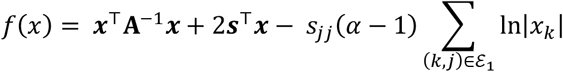

and a non-smooth term

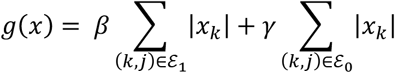

The variable *x* was iteratively updated as:

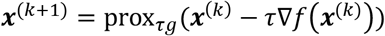

where *τ* denotes the step size. For the non-smooth part, the proximal operator was computed element-wise using the soft-thresholding function. The gradient of the logarithmic term was computed based on ∇_***x***_[−∑ ln|***x***|] = −***x***^−1^. To accelerate convergence, we adopted a FISTA scheme incorporating Nesterov momentum^41^. The algorithm was terminated when the relative change in the objective function in (Eq. 4) fell below 10^−5^ or when the maximum number of iterations (5000) was reached. For the PRECISE1K dataset, a threshold of 10^−4^ was adopted instead of 10^−5^ for computational efficiency.

### wgLASSO

Given a sample covariance matrix **S** ∈ ℝ^*p*×*p*^, wgLASSO estimates the precision matrix **Θ** by solving the following optimization problem:

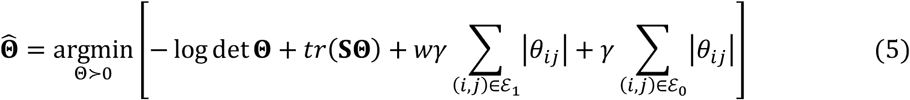

Here, *w* represents the relative weighting of sparsity applied to entries corresponding to known regulatory edges ℰ_1_. All other symbols are as defined in Eq. (1). The optimization of Eq. (5) was performed under the same conditions as in the gamma-guided graphical LASSO. The initial value of **Θ** was set to the solution obtained by the standard gLASSO. Runs in which the optimization became ill-conditioned or hyperparameter tuning failed (as defined below) were excluded from the analysis (Fig. 2D).

### GENIE3

GENIE3 infers gene regulatory interactions by modeling each gene as a function of all others using random forests, with variable importance defining edge scores^15^. Mean Decrease Impurity was adopted as the measure of variable importance. When estimating regulatory interactions to a known network, only random forest models with target genes within the network were constructed to reduce computational cost. For genome-wide network inference (Fig. 2B), random forest models were constructed for all genes, and the resulting scores were symmetrized by averaging (*i, j*) and (*j, i*). The sign of the inferred regulation (activation or repression) was determined by the sign of the Pearson correlation coefficient.

### Semi-supervised support vector machine (S3VM)

S3VM infers gene regulatory interactions by classifying gene pairs using a linear support vector machine trained on known regulatory edges. For the DREAM4 dataset, all remaining gene pairs were treated as negative samples. For the PRECISE-1K dataset, due to severe class imbalance, negative samples were randomly subsampled from remaining gene pairs to achieve a positive-to-negative ratio of 1:100. Feature vectors were constructed by the Hadamard product of the expression profiles of gene pairs. Alternative representations, such as concatenation, were evaluated but resulted in lower performance. The Kronecker product was not considered due to its excessive dimensionality. Separate SVM models were trained for activation and repression. The final regulatory score was defined as the decision value with the larger absolute value between the two models.

### Hyperparameter tuning

Hyperparameters were optimized by partially masking known regulatory interactions and evaluating their recovery performance. Specifically, known genes were randomly partitioned into five subsets. In each iteration, one subset was masked, and network inference was performed using the remaining known genes together with unknown genes. The estimated scores were ranked, and the ranks of the masked regulatory edges were extracted from this ranking. Scores equal to zero were assigned a rank of zero. This procedure was repeated for all subsets, and the sum of the ranks of the masked edges was used as the evaluation metric. Cases in which the inferred regulatory sign did not match the prior information were treated as failures and were not included in the rank-sum calculation. For gNAISTO, ranking during hyperparameter tuning was performed using 20 unknown genes to reduce computational cost. For wgLASSO, 500 unknown genes were used for ranking in the PRECISE-1K dataset, whereas all unknown genes were used for the DREAM4 dataset.

The hyperparameters α and β of the gamma distribution governing known edges in gNAISTO were determined by scaling reference values derived from the expression data. Specifically, the mean of the naive precision matrix entries corresponding to known edges 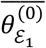 was computed as

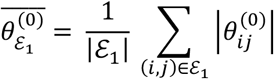

Here, 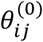 denotes an element of the naive precision estimate Θ^(0)^, which is defined as the inverse of the sample covariance matrix computed using only genes within the known network, rather than the full set of genes, to avoid instability in matrix inversion. The hyperparameters *α* and *β* were not directly optimized; instead, they were determined such that the mode (*α* − 1)/*β* and the ratio of variance to mean 1/*β* of the gamma distribution were scaled by 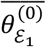. Specifically, the mode was varied as 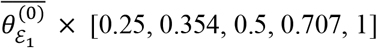, and the ratio of variance to mean as 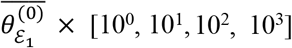. The hyperparameter γ was varied over [10^-6^, 10^-5^, 10^-4^, 10^-3^] for both gNAISTO and wgLASSO. For these methods, if the evaluation metric (rank-sum) was zero, the corresponding configuration was considered to reflect excessive regularization on unknown interactions, and all configurations with that γ were excluded from further exploration. For wgLASSO, the weight parameter *w* was varied over [1, 0.75, 0.5, 0.25]. For S3VM, the regularization parameter *C* was varied over [0.0675, 0.125, 0.25, 0.5, 1, 2]. No hyperparameter tuning was performed for GENIE3.

For wgLASSO applied to the PRECISE-1K dataset, the large number of variables led to substantial computational cost. To mitigate this, the regularization parameter *γ* was varied over relatively large values [10^-3.9^, 10^-3.6^, 10^-3.3^, 10^-3^]) to avoid ill-conditioned precision matrix estimation, and the number of unknown genes was restricted to 500 during hyperparameter tuning. If the evaluation metric (rank-sum) was zero for all configurations, hyperparameter tuning for wgLASSO was considered to have failed.

### Datasets

To evaluate inference performance, we used datasets from the DREAM4 in silico network challenge, a widely used benchmark for gene regulatory network (GRN) inference^4,17,18^. The datasets were downloaded from the GeneNetWeaver website (https://gnw.sourceforge.net/dreamchallenge.html). The benchmark consists of five distinct networks, each comprising 100 genes, together with simulated expression data generated using GeneNetWeaver. For each network, expression data from wild-type, knockout, and knockdown conditions (201 conditions in total) were merged and used for analysis. Datasets with measurement noise were generated by introducing synthetic noise into the noise-free data. To reproduce the mix of normal and log-normal noise, multiplicative noise (σ = 0.05) and additive noise (σ = 0.0001) were introduced as the baseline noise level (×1). This baseline was consistent with the characteristics of the original noisy dataset. We further validated our approach using PRECISE-1K, a large-scale curated *E. coli* expression dataset comprising 1,035 samples collected under diverse experimental conditions, including varying temperature, pH, and carbon sources^19^. From the provided transcriptional regulatory network, interactions for which regulator locus IDs could not be assigned or regulatory effects were not annotated as activation (+) or repression (−) were excluded, resulting in a network of 7,294 regulatory interactions. Genes with no connections in the network were removed, yielding a final set of 2,199 genes. Expression values were normalized per gene using z-score normalization prior to analysis. The known networks were sampled from the largest connected component of the ground-truth network using a random walk.

### Performance evaluation of network inference

The performance of the inferred regulatory network 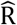 was evaluated against the ground-truth regulatory matrix R_true_ using the area under the precision–recall curve (AUPRC). In this study, both the presence/absence of edges and their regulatory sign (activation or repression) were considered. Specifically, an inferred edge was counted as a true positive only if it existed in the ground-truth network and its sign was correctly inferred; otherwise, it was counted as a false positive. The AUPRC was computed by integrating precision with respect to recall over varying decision thresholds.

## Code availability

The source code is publicly available on GitHub under the MIT License. The gNAISTO software package is available at https://github.com/sakulab-org/gNAISTO, and the analysis scripts used to generate the results presented in this study are available at https://github.com/sakulab-org/gNAISTO-paper. The gNAISTO repository includes installation instructions and example scripts demonstrating its usage.

## Acknowledgement

We thank members of our laboratory for discussion and feedback on this research. This work was supported by JSPS KAKENHI (JP25K18442 to TK; JP25K24212 to KTS; JP26K15049 to KK) and by Daiichi-Sankyo “Habataku” Support Program for the Next Generation of Researchers, NAIST Senju Monju Project. T.K and Y.S conceived and designed the project. T.K and Y.S developed the algorithm. T.K performed the computational evaluation. T.K., K.T.S., K.K, and Y.S wrote and revised the manuscript.

## Supplementary Information

**Fig. S1.**
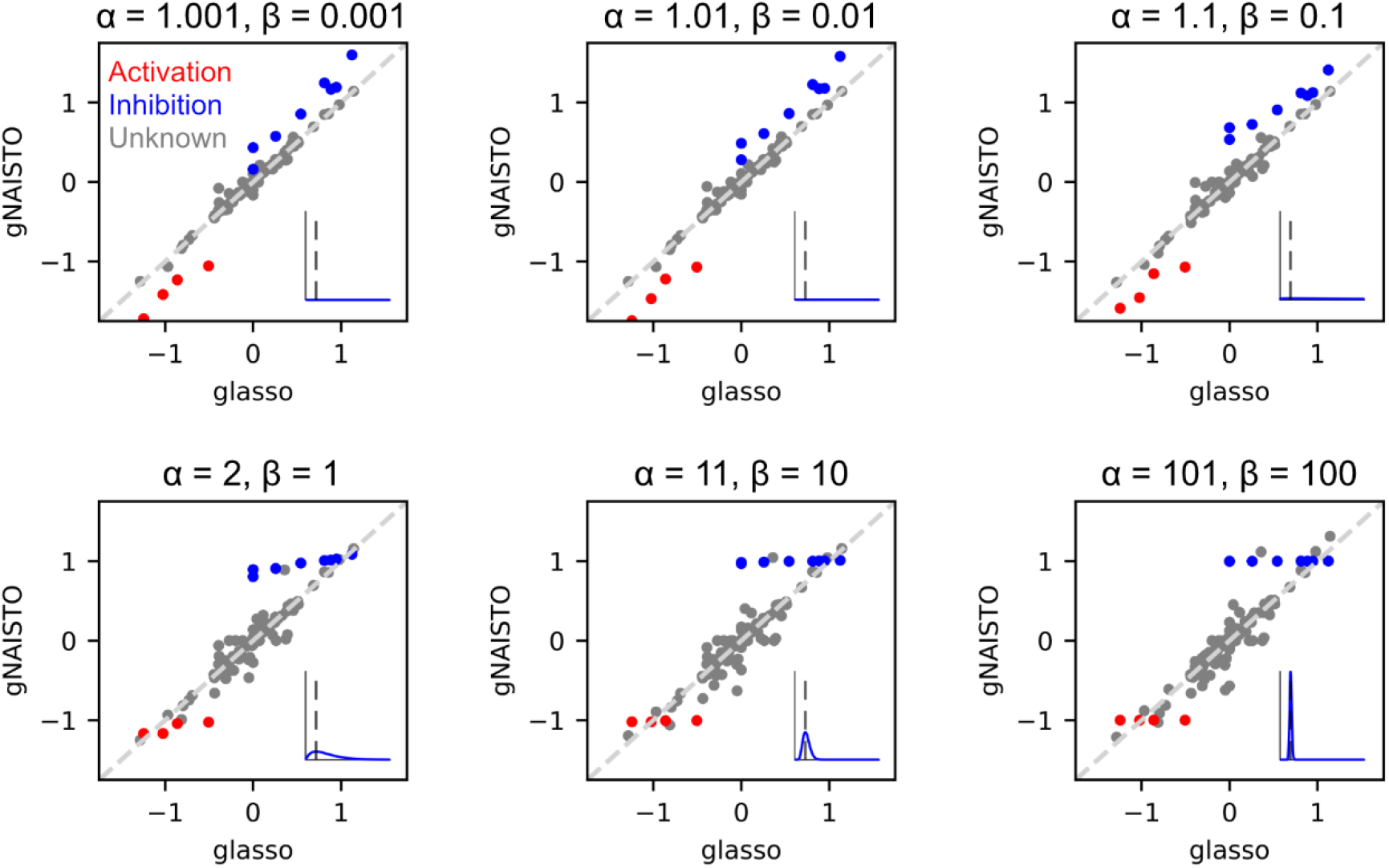
Comparison of precision matrices estimated by gNAISTO and gLASSO. Scatter plot comparing elements of the precision matrix estimated by gLASSO (x-axis) and gNAISTO (y-axis). A ground-truth precision matrix was generated from a randomly constructed scale-free network with 20 nodes, and simulated data were sampled from a multivariate normal distribution with the corresponding covariance matrix. Eight nodes and their connecting edges were designated as the known network. Elements corresponding to known edges are highlighted (red: activation; blue: repression). Note that edges are defined by the partial correlation coefficient 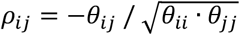, meaning that a negative *θ*_*ii*_ corresponds to an activating edge, while a positive *θ*_*ii*_ represents an inhibitory edge. The hyperparameters *α* and *β* of gNAISTO were chosen such that the mode of the gamma distribution satisfies (*α*−1)/*β*=1 (as illustrated in the inset). The hyperparameter *γ* was fixed at 0.05.

**Fig. S2.**
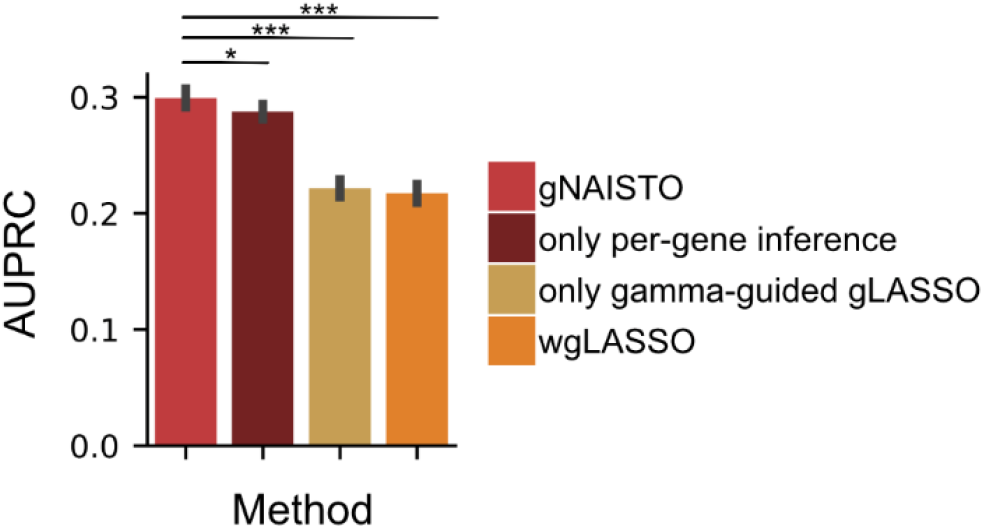
Impact of per-gene inference and gamma-guided gLASSO on AUPRC. Mean AUPRC values for the inference of regulatory edges between known and unknown genes using simulated datasets from the DREAM4 benchmark. Known networks were defined by randomly selecting 20 genes from the ground-truth networks. Error bars indicate the standard error across 50 trials (5 datasets × 10 random subnetworks). Statistical significance was assessed using the Wilcoxon signed-rank test (\**P* < 0.05, \*\**P* < 0.01, \*\*\**P* < 0.001).

